# The protamines of the noble false widow spider *Steatoda nobilis* provide an example of liquid-liquid phase separation chromatin transitions during spermiogenesis

**DOI:** 10.1101/2024.06.04.597381

**Authors:** Melissa R. Leyden, Peter Michalik, Luciana Baruffaldi, Susheen Mahmood, Ladan Kalani, Donald F. Hunt, Jose Maria Eirin-Lopez, Maydianne C.B. Andrade, Jeffrey Shabanowitz, Juan Ausió

**Affiliations:** Department of Chemistry, University of Virginia, Charlottesville, Virginia 22904, USA; Zoologisches Institut und Museum, Universität Greifswald, Greifswald, Germany; Department of Biological Sciences, University of Toronto Scarborough, 1265 Military Trail, Toronto, ON, M1C 1A4, Canada; Department of Biochemistry and Microbiology, University of Victoria, Victoria BC V8W 2Y2, Canada; Environmental Epigenetics Laboratory, Institute of Environment, Florida International University, Miami, Florida, USA

**Keywords:** Sperm nuclear basic proteins (SNBPs), protamines, spider, mass spectrometry, liquid-liquid phase separation, phylogeny

## Abstract

While there is extensive information about sperm nuclear basic proteins (SNBP) in vertebrates, there is very little information about Arthropoda by comparison. This paper aims to contribute to filling this gap by analyzing these proteins in the sperm of the noble false widow spider *Steatoda nobilis* (Order Araneae, Family Theridiidae). To this end, we have developed a protein extraction method that allows the extraction of cysteine-containing protamines suitable for the preparation and analysis of SNBPs from samples where the amount of starting tissue material is limited. We carried out top-down mass spectrometry sequencing and molecular phylogenetic analyses to characterize the protamines of *S. nobilis* and other spiders. We also used electron microscopy to analyze the chromatin organization of the sperm, and we found it to exhibit liquid-liquid phase spinodal decomposition during the late stages of spermiogenesis. These studies further our knowledge of the distribution of SNBPs within the animal kingdom and provide additional support for a proposed evolutionary origin of many protamines from a histone H1 (H5) replication-independent precursor.

## INTRODUCTION

Sexual reproduction in metazoan organisms involves two highly specialized cell types (gametes: sperm and oocytes) with important functional and epigenetic differences from their germ cell progenitors (Feng and Chen, 2015; Kota and Feil, 2010). Sperm exhibits significant structural (Baccetti and Afzelius, 1976; Pitnick et al., 2009) and chromosomal protein composition characteristics (Eirin-Lopez and Ausio, 2009), which involves three main groups of sperm nuclear basic proteins (SNBPs): H, histone; PL, protamine-like and P protamine types (Ausió, 1999). Histones (H-type proteins) have a composition rich in both lysine and arginine, vary in their mass between 10 and 25 kDa, and are the constituents of nucleosomes, the basic subunit of somatic chromatin (van Holde, 1988). During spermiogenesis, SNBPs of the PL and P types replace the somatic histones of the sperm progenitor cells that are present at the onset of spermatogenesis (Govin et al., 2004; Moritz and Hammoud, 2022; Oliva and H. Dixon, 1991; Rathke et al., 2014). Protamine-like proteins (Lewis and Ausió, 2002) are also rich in arginine and lysine. Still, they exhibit much more structural variability than histones and can range in mass between 6 and 40 kDa. Finally, protamines (Kasinsky et al., 2012) are highly arginine-rich. They can have up to 80% arginine, as in California market squid (*Loligo opalescens*) (Lewis et al., 2004a), ranging in mass from 3 to 10 kDa.

SNBPs have been characterized in the phylum Porifera (Ausió et al., 1997) and the classes Schiphozoa (jellyfish *Aurelia aurita, and Thaumatoscypha hexaradiatus*), Hydrozoa (*Catablema sp.; Mitrocoma cellularia* (Rocchini et al., 1996), *Hydractinia echinata* (Torok et al., 2023)) and Anthozoa [anemones, *Urticina (=Tealia) crassicornis*, *Anthopleura xanthogrammica* and *Metridium senile*] of the phylum Cnidaria (Rocchini et al., 1996) (Rocchini et al., 1995b). SNBPs have also been analyzed in the phylum Annelida (*Chaetopterus varipedatus*) (Fioretti et al., 2012) and quite extensively in the phylum Mollusca (Casas et al., 1993; Subirana, 1973) and in the phylum Chordata (Lewis et al., 2003; Oliva and Dixon, 1991; Saperas and Ausio, 2013). However, despite arthropods representing 80 % of all animals (Zhang, 2011) and the Order Araneae consisting of approximately 48,500 species (Dimitrov and Hormiga, 2021), information about the SNBPs in the phylum Arthropoda has been very limited (Leyden et al., 2024).

Within the SNBP types, the replacement of histones by protamines in vertebrate taxa was initially hypothesized to fulfill two main functions. 1) compaction of the DNA to streamline sperm mobility and to protect against damage during its journey in search of the egg and 2) to assist the erasure of transcriptional and epigenetic marks (Oliva and Dixon, 1991). However, as variation in the H, PL, and P SNBP types across the Tree of Life shows (Bloch, 1969; Eirin-Lopez and Ausio, 2009), compaction of DNA is not essential for fertilization. Nevertheless, Kasinsky hypothesized that internal fertilization might have constrained the range of SNBPs to protamines in amniotes (Kasinsky, 1989, 1995). Also, protamine amino acid composition, particularly arginine content, has been shown to affect sperm head shape and possibly involvement in sperm competition for the mammalian P1 protamine (Luke et al., 2016). In the functional regard, protamines have recently gained a lot of attention. In the case of invertebrate protamines, the protamine-mediated removal of histones in the sperm of *Drosophila* (Rathke et al., 2014) has been shown to protect paternal chromosomes from premature division at fertilization (Dubruille et al., 2023). In mice, it has been shown that despite these protamines’ seemingly monotonous arginine composition, substituting a single lysine in protamine 1 in this vertebrate results in sperm chromatin and reproductive fitness alterations (Moritz et al., 2023).

In this paper, we report the first-time characterization of the SNBPs of a spider, the noble false widow species *S. nobilis* (Araneae, Theridiidae). We show that the SNBPs correspond to the protamine type, discuss their occurrence in the context of arthropod phylogeny, and analyze their amino acid sequences. We also describe their involvement in spermatogenic chromatin condensation mediated by the spinodal decomposition (SD) and nucleation (Nc) dynamic mechanisms of liquid-liquid phase separation (LLPS) (Harrison et al., 2005; Kasinsky et al., 2021).

## Results

### SNBPs of *S. nobilis* within the context of arthropod phylogeny

Cysteine sporadically occurs in several SNBPs and has been described to be present in the protamines and other SNBPs of marine invertebrates (Giménez-Bonafé et al., 2002; Zhang et al., 1999), insects and vertebrates (Gusse and Chevaillier, 1978; Retief et al., 1995), where it is ubiquitously present in eutherian mammals (Balhorn, 2007; Oliva and H. Dixon, 1991).

Therefore, given our access to only a limited amount of material, we modified our SNBP extraction method (Leyden et al., 2024) to add a cysteine alkylation by pyridyl ethylation before the HCl protein solubilization, and we optimized our methods for small amounts of starting material (see Materials and Methods).

Figure 1A shows the electrophoretic pattern of the proteins extracted from the sperm from 54 *S. nobilis* pedipalps (SN), using this method, in comparison to the SNBP protamines from the California mussel (MC) and salmon (SL) as well as the histones from chicken erythrocytes (CM) that are used here as markers for protamines and somatic histones respectively. The former provides a relative estimate of the molecular mass (Mr) associated with the electrophoretic mobility, as in AU-PAGE, in contrast to SDS-PAGE, proteins do not run corresponding to their molecular mass. The electrophoretic pattern of *S. nobilis* SNBPS observed in this figure displays a complex mixture of bands with two predominant groups with relatively low and high electrophoretic mobility within the areas corresponding to histones and protamines. The former presumably arises from the contamination with the somatic tissue from the male pedipalp structures sampled here (in spiders, the sperm is stored in pedipalps before copulation (Zhang et al., 2022). Within the protamine region, an intense double band (Fig. 1A SN/P) exhibits an electrophoretic mobility similar to that of the PL-IV SNBP of the mussel *Mytilus californianus* (Fig. 1A, MC), which has a relative molecular mass (Mr) of 6.5 kDa (Carlos et al., 1993a).

**Fig. 1.**
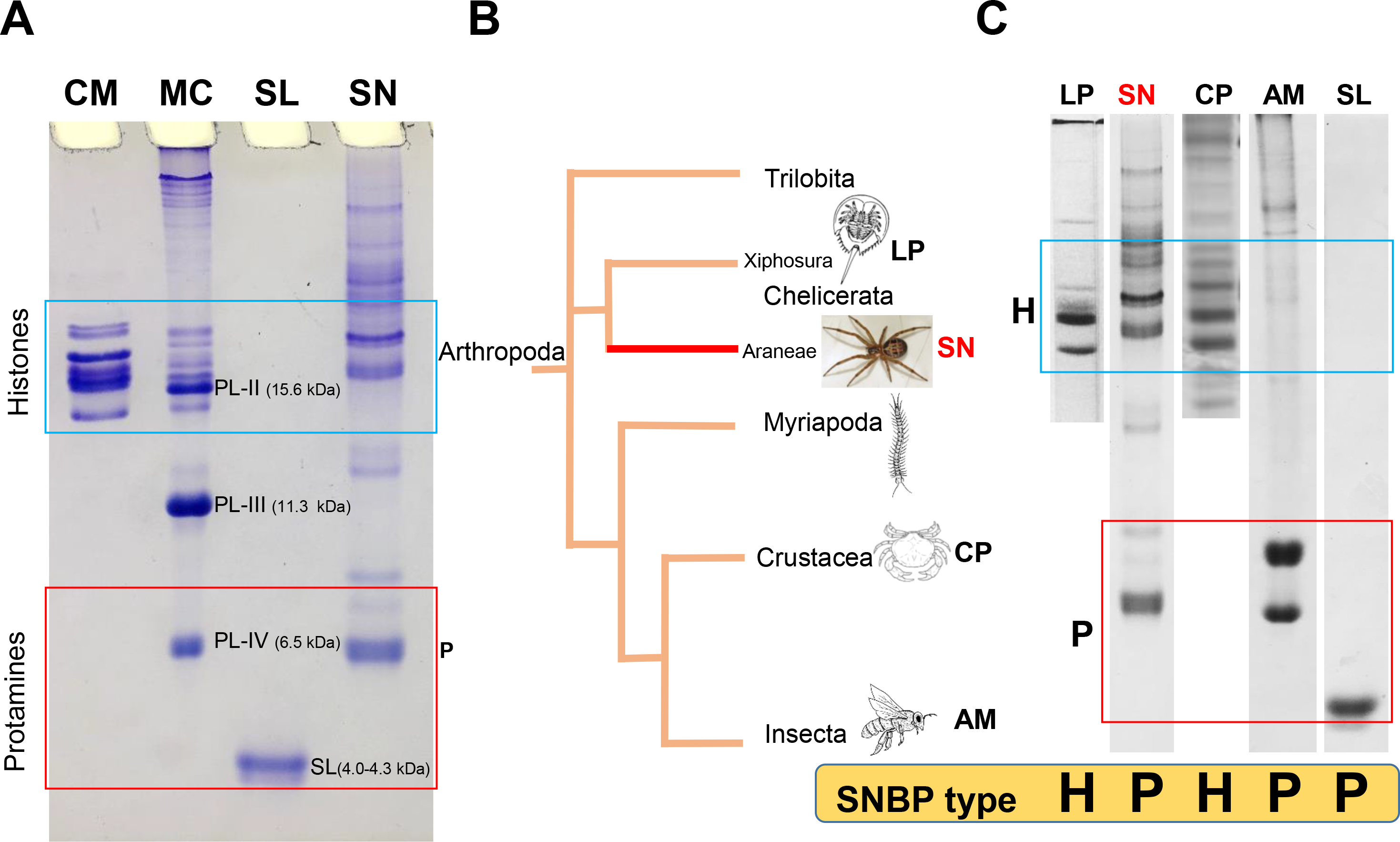
SNBPs of *S. nobilis* in comparison to other groups of arthropods. (A) Acid acetic urea PAGE of *S. nobilis* SNBPs (SN) in relation to the chromosomal proteins of other organisms used as protein markers. CM: chicken erythrocyte nuclear histones. MC: *Mytilus californianus* (California mussel) SNBPs indicating its major PL-II (Mr = 15.6 kDa) (Carlos et al., 1993b); PL- III (11.3 kDa) (Rocchini et al., 1995a) and PL-IV (6.5 kDa) components (Carlos et al., 1993a) SL: salmine protamine (4.0-4.3 kDa) (Ando, 1973) from the salmon *Onchorhyncus keta*. (B) Simplified phylogeny of Arthropods adapted from Giribet and Edgecombe (2019) (Giribet and Edgecombe, 2019). (C) Electrophoretic analysis of the SNBPs of several Arthropod representative species. LP: *Limulus polyphemus* (horse shoe crab) (Munoz Guerra et al., 1982); SN: *S. nobilis*; CP: *Cancer pagurus* (brown crab) (Kurtz et al., 2008); AM: *Apis mellifera* (honey bee) (unpublished) in comparison to salmine. H: Histones; P: Protamines.

Although this work analyzes only one spider species, and despite the large number of species and high complexity of the Order Araneae (Kulkarni et al., 2023; Lipke and Michalik, 2015), the occurrence of protamines in spiders documented here is of interest in terms of the SNBP phylogenetic distribution in the Phylum Arthropoda (Fig. 1B). Unfortunately, the amount of information available for this Phylum, which encompasses by far the largest number of existing species within the animal kingdom, is minimal. Figure 1C summarizes some of this information. Within the Order Xiphosura, the only SNBP information available is that of the horseshoe crab *Limulus polyphemus,* which, as shown in (Fig. 1C, LP) belongs to the H type.

This is similar to what is observed in the brown crab *Cancer pagurus* (Fig. 1C, CP) (Kurtz et al., 2008), which appears to be a common occurrence in decapod crustaceans (shrimp and crabs) (Chen et al., 2019; Wu et al., 2015). In insects, a protamine-like SNBP has been described in *Drosophila* (Jayaramaiah Raja and Renkawitz-Pohl, 2005), and a protamine is present in the sperm of the honey bee (*Apis mellifera*) [Fig. 1C AM (Leyden et al., 2024)]. Therefore, the information available so far suggests that as in vertebrates (Eirin-Lopez and Ausio, 2009) and plants (Borg and Berger, 2015; D’Ippolito et al., 2019), SNBPs are sporadically distributed in arthropods and that a transition from the H to the P SNBP types might have occurred several times (Ausió, 1999) throughout the evolution of this group.

### Characterization of *S. nobilis* protamines

We recently published the amino acid sequences of SNBPs from several insect species (Leyden et al., 2024). Following a similar mass spectrometry (MS) approach as that described in (Leyden et al., 2024), two partial closely related sequences of proteins (P1, P2) with molecular masses 6538.8 and 6395.9 were obtained (Fig. 2A) corresponding to the protamines shown in Fig. 1A. A genome BLAST using these sequences identified a hypothetical histone H1-H5 protein AVEN_19411.1 from *Araneus ventricosus* (Family Araneidae) (Kono et al., 2019) (Fig. 2B-1), which encompassed a highly homologous protein of a very similar size (Mr ∼ 6200) at its C-terminal domain (Fig. 2B 1-2). Further searching using the *A. ventricosus* sequence allowed us to obtain an alignment of more spider sequences (Arakawa et al., 2022) (Fig. 2C) and identify a putative consensus protamine sequence for spiders of the Araneidae and Theridiidae families (Fig. 2D). Unfortunately, we were not able to retrieve sequences from any other Families. It is interesting to note that the two partial P1 and P2 amino acid sequences from *S. nobilis* (Fig. 2A) indicate that, as in insects (Leyden et al., 2024), the spider protamines also exhibit sequence microheterogeneity, likely as a result of the existence of more than one encoding gene. The isoelectric point of all these proteins (12.2 – 12.5) is well within the range observed for insect protamines.

**Fig. 2.**
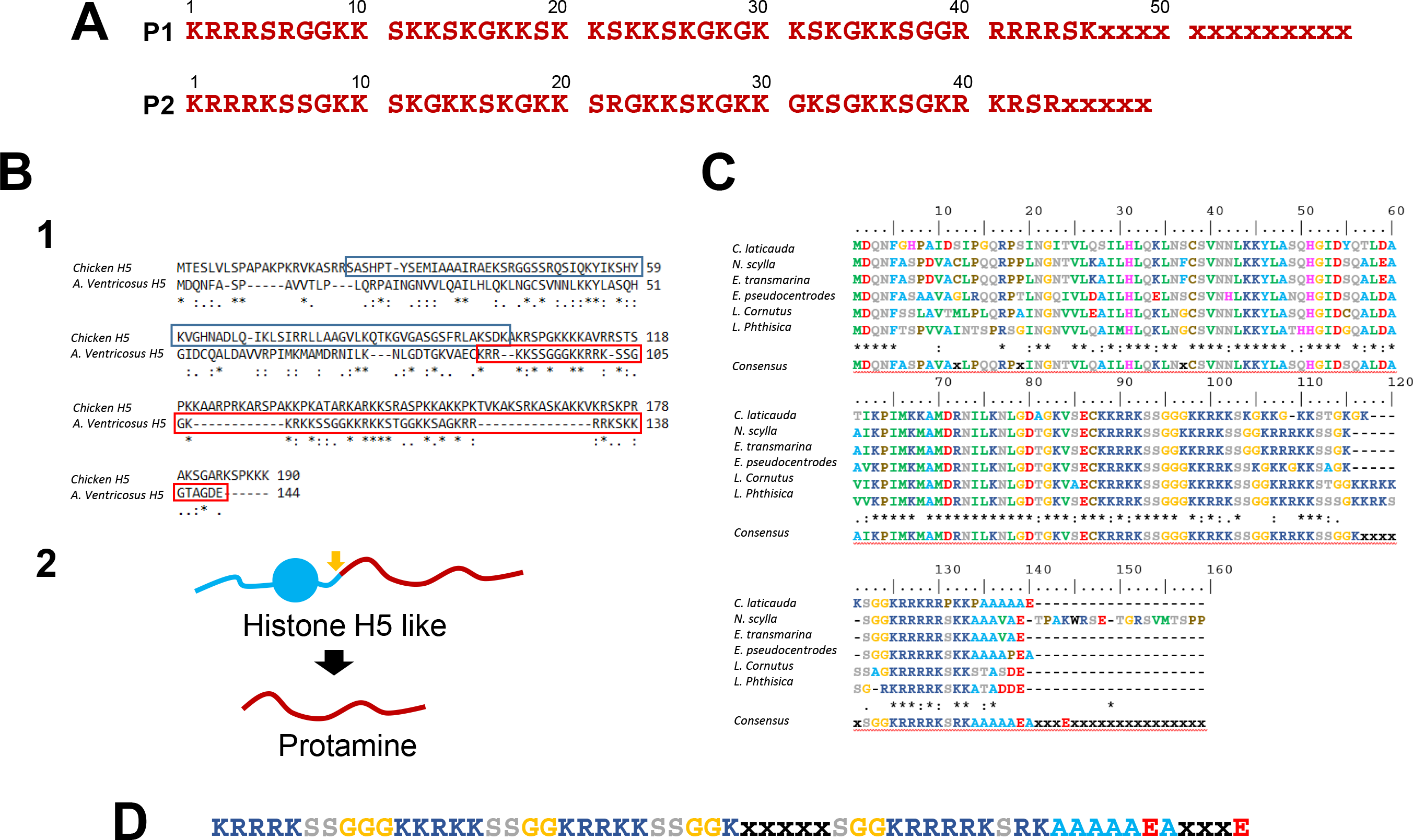
Protein sequences of spider protamines. (A) Partial amino acid sequences of the P1 (∼6359Da) and P2 (∼6396Da) protamines from *S. nobilis* determined by LC-MS. (B) 1. Amino acid sequence alignment of chicken histone H5 (AAA48798.1) and *A. ventricosus* H1-H5 like protein (GBL99906). 2. The protamine region (red) is part of the C-terminal region of a histone H5-lke protein (blue). (C) Sequence alignments of the histone H5-like proteins from different spiders: *Cyclosa laticauda* (IAWK01030102.1); *Neoscona scylla* (ICBS01001118.1); *Eriophora transmarina* (IBPW01032809.1); *Eriovixia pseudocentrodes* (ICEC01024091.1); *Larinoides cornutus* (IBUI01019351.1) and *Larinia phthisica* (IAJV01034118.1). In these alignments, (*) indicates that the amino acid is the same for all the sequences at that position and (:) indicates that some of the sequences have different amino acids at that position, but that the chemical properties of the different amino acids are similar. (D) Consensus sequence of spider protamines.

Of interest, in addition to the protamine sequences, the high sensitivity of our MS analyses allowed us to determine the presence in our sample (Fig. 1A) of peptides such as: DEKKKDDKKSGTGKPQQKPEEKKPEKGGKKDEKKPEKKPEQKKK (∼5101Da), MKEKPDDKGKPGEKKPEGPKPGEKKPEPGKPGEKKPGEPKPGEKKPEEKGPKK (∼5810Da) and MDKKPDDKGKPGEKKPELGKPGEKKPDDKggKPGEKKPEEKGPKK (∼4953Da). In the last two instances, the sequences contained oxidized methionines and acetylated N-termini, with the lowercase residues not confirmed. A BLAST of these proteins against the NR database (with no organism chosen) revealed some extent of homology to proteins of bacterial origin. This is not surprising as the sperm used in our analysis was obtained from spider pedipalps (see Materials and Methods) and contained a significant amount of bacteria, as visualized by EM (results not shown). The relevance of this to our protamine analysis will be discussed below.

### Liquid-liquid phase separation chromatin transition during *S. nobilis* spermiogenesis

To visualize the process of spermiogenesis in *S. nobilis*, electron microscopy was performed of *Steatoda grossa*, a closely related species. Figure 3 shows the ultrastructural changes the nucleus and chromatin undergo during the differentiation process. Interestingly, late spermatids undergo spinodal decomposition (SD) (Fig. 2 a-b) and nucleation chromatin condensation (Fig. 2c-d) liquid-liquid phase separation (LLPS) transitions. This is similar to what it was initially described for the mollusc *Murex brandaris* (Harrison et al., 2005) and to what has been extensively observed during insect spermiogenesis (Kasinsky et al., 2021). Yet, this is far from being an invertebrate chromatin condensation phenomenon; previous evidence indicates this also takes place in condrychtian fish spermiogenesis (Gusse and Chevaillier, 1978), and somatic chromatin has also been described to involve LLPS (Gibson et al., 2019).

**Fig. 3.**
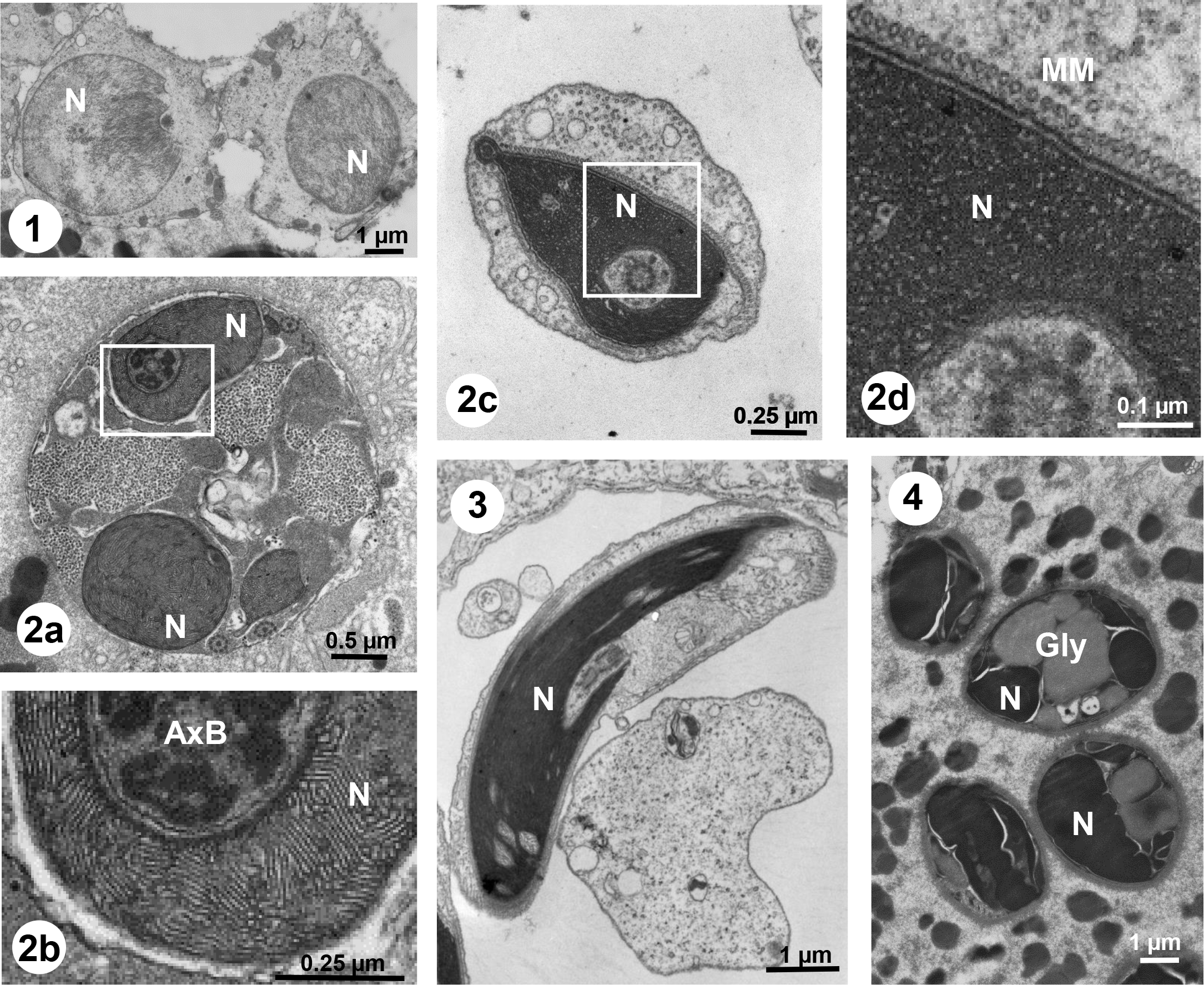
Chromatin transitions during *Steatoda grossa* spermiogenesis. (1) Early spermatids with beginning of chromatin condensation; (2 a-d) Cross-sections of late spermatids at different stages of chromatin condensation undergoing spinodal decomposition (SD) (2 a-b) and nucleation (2 c-d). (3) Longitudinal section of a late spermatid. (4) Mature sperm. AxB, axonemal basis; Gly, glycogen; N, Nucleus; MM, manchette of microtubules.

Analysis of the pattern in Fig. 3-2b yielded a value of 170 ± 20 Å for the distance between the chromatin lamellae [which corresponds to the lambda parameter used in the SD calculations (Harrison et al., 2005)] and a chromatin lamellae thickness of approximately 50 Å (Fig. 4A-3).

**Fig. 4.**
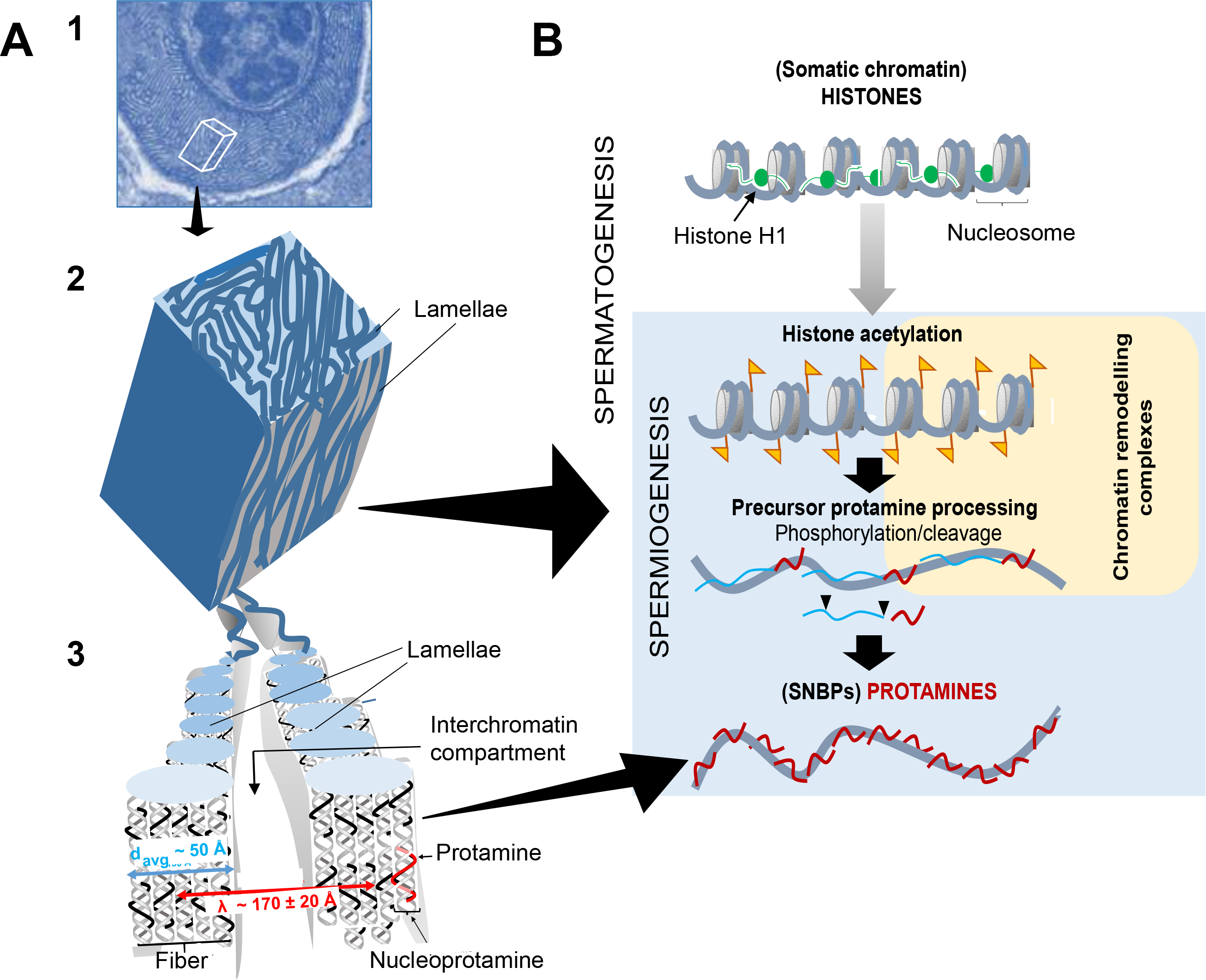
Chromatin transitions during *S. nobilis* spermatogenesis. (A-1) Fig. 3-2b showing spermatid chromatin undergoing spinodal decomposition. (A-2) Three D schematic representation of the cube section indicated in (A-1). (A-3) Representation of the dynamic lamellae in (A-2). (B) Cartoon representation of the major chromatin changes during *S. nobilis* spermatogenesis. During post-meiotic spermiogenesis, histones of the nucleosomal chromatin fiber become hyperacetylated (yellow flags) in preparation for histone eviction and replacement by protamine precursors. This is a highly dynamic complex process that involves histone degradation and several precursor protamine post-translational modifications (cleavage and phosphorylation) presumably assisted by chromatin remodeling complexes and chaperone proteins that takes place at the interchromatin compartment (Cremer et al., 2020). By the late stages of spermiogenesis, protamines replace most of the germ cell progenitors.

The chromatin in the mature sperm found in the spermatheca is completely condensed, and glycogen is present in substantial amounts (Fig.3-4). The coexistence of glycogen with mature sperm in spiders has been well documented (Coyle et al., 1983; Michalik et al., 2004; Michalik et al., 2005). The presence of glycogen is quite widespread in both invertebrate and vertebrate species (Anderson and Personne, 1970) and although its function is still controversial, it appears to have an important yet overlooked contribution to different aspects of spermatogenesis (Silva et al., 2022).

## DISCUSSION

### Of clams, chickens, and spiders

An early of the SNBPs from molluscs revealed extensive protein heterogeneity both in size and composition between the species analyzed (Subirana, 1973), with those with smaller molecular mass generally exhibiting a higher arginine content. A hypothetical evolutionary pathway was proposed at the time in which the arginine-rich protamine-like proteins and protamines might have been derived from a ’strongly basic fragment’ of a histone precursor (Subirana, 1973). However, the precise molecular details remain unknown. When in 1987 we determined the amino acid sequence of the globular domain of the protamine-like (PL) protein from the sperm of the surf clam *Spisula solidissima*, a high extent of homology was found with that of the winged helix domain (WHD) (Ramakrishnan et al., 1993) of the histone H5 of chicken erythrocytes (Ausió et al., 1987). Histone H5 is a replication-independent member of the histone H1.0 family of histone H1 linker proteins, which are often present in terminally differentiated cells (Cheema and Ausio, 2015), such as the nucleated erythrocytes of fish, amphibians reptiles and birds (Shay et al., 1988). Such an observation provided the first hint that sperm protamines and PLs might have evolved from a common ancestral histone H1 (Ausió, 1995, 1999). Since then, information has been accumulating in support of this proposal in such diverse organisms as a marine annelid *Chaetopterus variopedatus* (Fioretti et al., 2012), tunicates (Lewis et al., 2004b; Saperas and Ausio, 2013) and more recently in the liverwort *Marchantia polymorpha* (D’Ippolito et al., 2019). The protamines of *Steatoda nobilis* and those of the superfamily Araneoidea (Fig. 2) provide further support to the histone H1 hypothesis (Ausió, 1999).

In most of the organisms where the final SNBP at the end of spermiogenesis was a protamine arising from a histone H1-H5, the precursor was clearly identified in the samples analysed. Yet, this is not the case here, despite the high sensitivity of the MS approach used. This may be because, in contrast to previous studies, in the case of *S. nobilis* only the protein products present in the mature sperm obtained from the spider pedipalps were used in the analysis. The detection of bacterial protein/peptides in this study is consistent with this, as spiders like *S. nobilis* use intromittent organs to transfer sperm to females, and environmental microbes are introduced into females during mating (Spicer et al., 2019).

### SNBP distribution in Arthropoda

The occurrence of a protamine-like arginine was described earlier using cytochemical staining (Rasch and Connelly, 2006), yet no information on the molecular details have been reported. The presence of protamines in the Order Aranea (Fig. 1 B-C) is interesting when visualized in the context of Arthropoda evolution. As in fishes (Saperas et al., 1997) and amphibians (Ausio et al., 2007) within the Phylum Chordata, the SNBPs of the Phylum Arthropoda also appear to exhibit a sporadic distribution (Saperas et al., 1994) (Fig. 1C).

Although it would be tempting to corelate the SNBP types to the living environment with the H type SNBP occurring in the aquatic environment versus the P type corresponding to the terrestrial species, this is unlikely to be the case. In this regard, the three main SNBP types (H, PL and P) are present in Fish. The same is also true if the internal versus external fertilization type is considered. While *Limulus polyphemus* (Fig. 1C, LP) has external fertilization (Brockmann et al., 1994), crustaceans have both external and internal fertilization (Aiken et al., 2004) but the crab *Cancer pagurus* (Fig. 1C, CP) has internal fertilization (Edwards, 1966). In contrast, it appears that at least in the spider *S. nobilis* and in insects such as in *Apis mellifera*, both with internal fertilization, protamines constitute the prevalent type (Fig. 1C, SN, AM).

Hence, as Chordata, the SNBP types in Arthropoda also exhibit a sporadic distribution, and in both instances, the evolutionary cause remains unclear. As we had earlier proposed, such heterogeneous distribution could be possibly explained by a repeated and independent loss of the expression of the protamine gene (or loss of the gene itself) (Saperas et al., 1994) in the course of Metazoan evolution.

### Ubiquitous occurrence of LLPS by spinodal decomposition during metazoan spermiogenesis

When the mechanism of protamine-driven LLPS underlying sperm chromatin condensation by spinodal decomposition and nucleation was initially described (Harrison et al., 2005), it might have appeared to be a rare phenomenon sporadically distributed. However, upon closer inspection, it shows persistent occurrence in insects (Kasinsky et al., 2021), and we show here that it also occurs in spiders, suggesting this mechanism is by far more widespread than originally anticipated.

One of the first detailed studies describing the lamellar organization, which is characteristic of spinodal decomposition during spermiogenesis, was carried out in the condrichthyan lesser spotted fish *Scylliorhinus canicula* (Gusse and Chevaillier, 1978). The authors were able to characterize the different protein transitions undergone during spermiogenesis taking advantage of the testicular zonal organization consisting of different cell type layers corresponding to the stages of differentiation (Gusse and Chevaillier, 1981). We now know that such peculiar chromatin lamellar organization is the result of a highly dynamic process of LLPS known as spinodal decomposition (SD) (Harrison, 2010; Harrison et al., 2005). This is a dynamic process that, at the biochemical level involves a massive degradation of most of the genomic histones and post-translational processing of both chromosomal proteins involved (histone acetylation and protamine phosphorylation and cleavage) (Fig.4). Hence, this process is not only highly dynamic but it is also very complex (Kasinsky et al., 2012). The difficulty in the study of the biochemical events involved in the chromatin transitions at the different stages of spermiogenesis, particularly in invertebrates, where only low amounts of sample are usually available, arises from the difficulty in fractionating the different cell types at each stage.

Whether additional proteins are involved, such as the coiled-coil glutamate-rich protein 1 (CCER1), recently described in mice that participate in the histone to protamine transition during post-meiotic spermatid differentiation through formation of a liquid-liquid phase-separated condensate (Qin et al., 2023), is not known. Regardless, all these biochemical events leading to the particular chromatin patterning observed in SD (Fig. 4) occur through reactions that take place in the interchromatin compartment (Cremer et al., 2020), which is the main player in the different chromatin transitions observed.

Little is known about lamellar nucleoprotamine organization (Fig. 4A-3). Whereas chromosomal proteins that bind to DNA in a sequence-specific way preferentially bind to the major groove (Garvie and Wolberger, 2001) those that bind in a non-sequence specific way, such as histones (Luger et al., 1997), usually do it through interactions with the minor groove (Murphy and Churchill, 2000). Despite this, it has long been known that protamines, which also bind DNA with non-sequence specificity, bind to the major groove (Fita et al., 1983; Hud et al., 1994; Mukherjee et al., 2021; Puigjaner et al., 1986). Recent work solved the apparent puzzle by showing that phosphorylation reduces the binding affinity of the minor groove by introducing an important deformation that eliminates the minor groove preference of protamines (Chhetri et al., 2023). Hence, serine phosphorylation at the time of protamine deposition onto DNA during the replacement of histones by protamines (Balhorn, 2007; Oliva and Dixon, 1991; Willmitzer and Wagner, 1980) (Fig. 4B) appears to play a critical role in SD and in the resulting organization of the nucleoprotamine complexes along the lamellae (Fig. 4A-3). Importantly, *S. nobilis* protamines (Fig. 2A) contain several serines at positions similar to the RRRS amino acid motifs of the peptides used in the Chhetri study (Chhetri et al., 2023).

## MATERIALS AND METHODS

### Biological material

*Steatoda nobilis*, (Thorell, 1875) the noble false widow, is a cobweb building spider species in the Theridiidae family. It is an invasive and synanthropic species (Dunbar et al., 2020) native to the Canary Islands or Madeira that has been spread by human activity to almost every continent (Bauer et al., 2019). Females and males can be found living in urban areas around human-made structures (Dunbar et al., 2020). The *S. nobilis* males used in this experiment were the laboratory F1 offspring of field-mated females collected in 2022 from a population in Nottingham UK (Cullen, 2017). Males were grown in temperature and light-controlled rooms (21-26 °C, 12h:12h light:dark) at the Andrade Laboratory at the University of Toronto in Scarborough. Similar to other spider species (Zhang, 2011) *S. nobilis* males do not have a direct connection between their gonads (glands responsible for sperm production) and their paired intromittent organs (pedipalps, structures responsible for carrying and transferring the sperm during mating, (Foelix, 2011).

Therefore, after reaching adulthood, males eject sperm from their genital opening onto a small web and take it up into their pedipalps during a process called sperm induction (Foelix, 2011). In spiders, sperm cells are encapsulated with a protein sheath and can be transferred from the pedipalps to the female’s genital duct via a sclerotized structure on the palp (the embolus (Foelix, 2011).

### Sperm collection

Unmated adult males anesthetized under CO_2_ were transferred to a Petri dish under a dissecting microscope. Their pedipalps were removed using iridectomy microdissection scissors and placed in a LoBind 1.5 mL micro test-tube (Eppendorf) containing 40 µL of sperm solution: [220 mM NaCl, 10 mM CaCl_2_, 10 mM MgCl_2_, 7 mM KCl, 0.1% Triton X-100, Tris-HCl (pH 8.2)] (Modanu et al., 2013; Snow and Abndrade, 2004), and the pedipalps were crushed with a micropestle (Argos Technologies, Inc.) to release the encapsulated sperm cells.

Thereafter, another 35 µL of sperm solution was added to rinse the remaining sperm cells from the micropestle into the test-tube, for a total of 75 µL in each tube. The tube containing the sperm sample was then vortexed for approximately 30 sec and centrifuged at 1000 x g for 10 min, and this process was repeated a total of three times (Snow and Abndrade, 2004). After the last round of centrifugation, the tube was filled (close to their maximum volume) with 80% ethanol. The same procedure was conducted for a total of 54 adult unmated males. The samples were shipped to the University of Victoria.

### Protein extraction

Proteins were extracted from fifty-four pedipalps prepared as described above. At the University of Victoria, ethanol was removed by centrifugation (10 minutes at 4 LJC and 7,850 x g), and the pellet was vacuum dried and Dounce homogenized in 2 mL oh 0.6 N HCl. The homogenate was centrifuged in an Eppendorf microfuge at 16,000 x g, and the supernatant was precipitated with six volumes of acetone and overnight incubation at -20 LJC. Next day the sample was centrifuged at 7,850 x g and the pellet was vacuum dried and dissolved in 1,5 mL of ddsH_2_O.

### Method to extract cysteine-containing SNBPs from small amounts of sample using pyridylethylation

We use mouse testes to optimize a method for the extraction of cysteine-containing protamines.To this end, mouse testes were homogenized in buffer: 0.25 M sucrose, 60 KCl, 15 mM NaCl, 10 mM MES (pH 6.5), 5mM MgCl2, 1mM CaCl2, 0.5% triton X-100 containing 1:100 protease inhibitor cocktail (Roche; 5056489001) using Dounce with ten strokes on ice. Approximately 1.5 mL of buffer per one testis was used. The purpose of this step was to remove the fat and part of the connective tissue. The homogenate thus obtained was centrifuged at 2,000 x g, and the pellet thus obtained will be referred to as mouse testes. Approximately 20 milligrams of mouse testes (∼15-20 milligrams of spider/insect testes) were re-suspended in 50 µL of 4M guanidinium chloride, 50 mM Tris-HCl (pH7.5), 1.25 mM EDTA and thoroughly vortexed.

Upon addition of 1 µL of β-mercaptoethanol, the sample was homogenized with a microcentrifuge polypropylene pellet pestle and incubated for 90 min. in the dark. Next, 2 µLs of vinyl pyridine were added to the mixture and the sample was further incubated in the dark with brief vortexing at 5 min. Intervals. After incubation, 400 µLs of 0.6 N HCl were added to the mixture and homogenized using a 2 mL Wheaton glass Dounce on ice with 10 strokes. The homogenate was centrifuged at 16,000 x g at 4 LJ C in an Eppendorf microfuge. The supernatant was split into two 220 µL aliquots (in Eppendorf tubes), precipitated by mixing it with 6 volumes of acetone, and let stand overnight at -20 LJC. The precipitate thus obtained was centrifuged at 16,000 x g at 4 LJC. The supernatant was discarded, and the pellet was speed vacuumed for 10 min. at room temperature and used for further analysis.

### Gel electrophoresis

Acetic acid (5%)–urea (2.5 M) polyacrylamide gel electrophoresis (AU-PAGE) was carried out according to Hurley (Hurley, 1977) and, as described elsewhere (Ausió, 1992).

### Reversed-phase HPLC (RP-HPLC)

HPLC was performed as described (Ausió and Moore, 1998; Cheema and Ausio, 2017). In brief, a 1,000 µL aqueous solution of the protein extract from the sperm of 54 HCl-extracted pedipalps were injected onto a C_18_ column (Vydac, Hesperia, CA, USA.) (4.6 x 250 mm, particle size: 5 µm, pore size: 300 Å) and eluted at 1 mL/min using a mobile phase consisting of (0.1% trifluoroacetic acid) and acetonitrile gradient. Samples were fractionated on a Beckman Coulter SYSTEM GOLD® 126 Solvent Module equipped with SYSTEM GOLD® 168 Detector.

### Electron microscopy

Male specimens were dissected in 0.1 M phosphate buffer (PB) to which 1.8 % sucrose was added. The reproductive system was fixed for 2 hours in 2.5 % glutaraldehyde in PB and postfixed for 2 hours in PB buffered 2 % OsO_4_. After being washed in PB, samples were dehydrated in graded ethanols and embedded in Spurr’s resin (Spurr, 1969). Ultrathin sections (60 nm) were obtained using a Diatome Ultra 45° diamond knife on a Leica ultramicrotome UCT. Sections were stained with uranyl acetate and lead citrate following Reynolds (Reynolds, 1963) and examined with a JEOL TEM 1011 electron microscope at 80 kV. Images were captured with an Olympus Mega View III digital camera using the iTEM software.

### Mass spectrometry

Pierce LC-MS grade water and formic acid were purchased from Thermo Scientific (Rockford, IL). LC-MS grade acetonitrile and 2-propanol were purchased from Honeywell (Charlotte, NC). Acetic acid was purchased from Sigma Aldrich (St. Louis, MO).

For hydrophilic interaction chromatography (HILIC) 1μL of extracted protein solution in 0.5% acetic acid was diluted 10-fold with acetonitrile. Approximately 1μL (95%) of the dilution, corresponding to ∼5% of the total suspension of the unfractionated samples, was pressure-loaded onto in-house prepared pre-columns (360μm OD x 100μm ID) (Udeshi et al., 2008).

Fractionated samples were reconstituted in 1% acetic acid in water and diluted with acetonitrile to 80% acetonitrile. 10% of the original fractionated solution were pressure loaded onto pre-columns. Both analytical and pre-columns had a 2mm Kasil 1624 frit, and the analytical column (360μm OD x 75μm ID) had a laser-pulled electrospray tip (Ficarro et al., 2009). The HILIC pre-column was packed to 7cm with 12μm diameter, 300Å PolyHYDROXYETHYL A (PHEA) packing material from PolyLC Inc. (Columbia, MD) and was connected to an analytical column packed to 10cm with 5μm diameter, 300Å PHEA packing material. For reverse-phase chromatography 5% of the extracted protein solution was pressure loaded onto a reverse-phase column in 0.5% acetic acid. The reverse-phase pre-column was packed to 7cm with 10μm diameter, 300Å PLRP-S packing material from Agilent (Santa Clara, CA) and connected to an analytical column packed to 10cm with 3μm diameter, 300Å PLRP-S packing material.

An Agilent Technologies (Santa Clara, CA) 1100 Series Binary HPLC system coupled to a Thermo Scientific Orbitrap Fusion Tribrid mass spectrometer (San Jose, CA) operated in low pressure Intact Protein Mode was used to analyze the proteins in each sample.

The PLRP-S pre-column was rinsed with 100% solvent A (0.3% formic acid in water) for 20 minutes at a flow rate of ∼3μL/min and then connected to the PLRP-S analytical column.

Proteins were eluted using a gradient of 0-60-100% solvent B (72% acetonitrile, 18% 2- propanol, 10% water, and 0.3% formic acid) in 0-60-70 minutes at a flow of ∼100nL/min. Highly basic proteins are not retained well on reverse-phase columns [3], and for this reason,

HILIC was used to retain the highly hydrophilic proteins in the samples. The HILIC PHEA- packed pre-column was washed with solvent B (95% acetonitrile, 15% water, 0.2% acetic acid) for 20 minutes at a flow rate of ∼3μl/min and then connected to a PHEA-packed analytical column. Proteins were eluted using a gradient of 100-0% solvent B for 60 minutes with a 10- minute hold of 100% solvent A (0.5% acetic acid in water) before re-equilibrating the column back to 100% solvent B at a flow rate of ∼100nL/min (Buszewski and Noga, 2012).

Proteins were selected for fragmentation from a 60,000 resolution Orbitrap MS1 scan. Using a 3-second cycle time, proteins with a charge state ≥3 were isolated by the quadrupole with an isolation window of 2m/z and fragmented by electron transfer dissociation (ETD) for 5ms and collisional dissociation (Compton et al., 2012; Mikesh et al., 2006). MS2 scans were acquired in the Orbitrap at 120,000 resolution with an automatic gain control target of 1e5.

MS1 and MS2 spectra were manually inspected using Qual Browser (Thermo Scientific).

MS2 ETD spectra were deconvolved using the Xtract algorithm (Thermo Scientific) (Senko et al., 1995). The protein sequences were determined by manual *de novo* analysis of MS2 spectra. Identified proteins were searched by BLAST to identify potential protein matches in bacterial and spider species (Boratyn et al., 2013).

## Acknowledgments

Thanks to Laini Taylor for collecting S. nobilis, Andrade lab assistants for help rearing the spiders, David Punzalan for introducing MCBA to JA and initiating the conversations that led to this study.

## Declaration of Interest

The authors declare that there is no conflict of interest that could be perceived as prejudicing the impartiality of the study reported.

## Funding

This work was funded by Natural Science and Engineering Research Council of Canada (NSERC), Grant RGPIN-2017-04162 to JA. *S. nobilis* collection, rearing and pedipalp preparation were funded by the Natural Sciences & Engineering Research Council of Canada (NSERC, Discovery Grant #2022-03597 to MCBA). DFH funding was from National Institutes of Health (NIH) grant GM037537.

## Author contribution statement

M.R.L. performed mass spectrometry; P.M., performed electron microscopy; L.B. and S.M. collected the sperm samples; L.K. was involved in the biochemical fractionation and processing; B.G. performed electron microscopy; M.C.B.A. conceptualization, funding and editing; D.F.H. and J.S. mass spectrometry; J.M. E-L. phylogenetic analyses; J.A. protein isolation, writing of the manuscript and conceptualization.

## Notes

### Competing Interest Statement

The authors have declared no competing interest.

